# CRISPR Spacers Acquired from Plasmids Primarily Target Backbone Genes, Making Them Valuable for Predicting Potential Hosts and Host Range

**DOI:** 10.1101/2023.12.07.570633

**Authors:** Lucy Androsiuk, Sivan Maane, Shay Tal

## Abstract

In recent years, there has been a surge in metagenomic studies focused on identifying plasmids in environmental samples. While these studies have unearthed numerous novel plasmids, enriching our understanding of their environmental roles, a significant gap remains: the scarcity of information regarding the bacterial hosts of these newly discovered plasmids. Furthermore, even when plasmids are identified within bacterial isolates, the reported host is typically limited to the original isolate, with no insight into alternative hosts or the plasmid’s potential host range. Given that plasmids depend on hosts for their existence, investigating plasmids without knowledge of potential hosts offers only a partial perspective.

This study introduces a method for identifying potential hosts and host ranges for plasmids through alignment with CRISPR spacers. To validate the method, we compared the PLSDB plasmids database with the CRISPR spacers database, yielding host predictions for 46% of the plasmids. When compared to reported hosts, our predictions achieved an 84% concordance at the family level and 99% concordance at the phylum level. Moreover, the method frequently identified multiple potential hosts for a plasmid, thereby enabling predictions of alternative hosts and the host range.

Notably, we found that CRISPR spacers predominantly target plasmid backbone genes while sparing functional genes, such as those linked to antibiotic resistance, aligning with our hypothesis that CRISPR spacers are acquired from plasmid-specific regions rather than insertion elements from diverse sources. Lastly, we illustrate the network of connections among different bacterial taxa through plasmids, revealing potential pathways for horizontal gene transfer.

**IMPORTANCE:** Plasmids are notorious for their role in distributing antibiotic resistance genes, but they may also carry and distribute other environmentally important genes. Since plasmids are not free-living entities and rely on host bacteria for survival and propagation, predicting their hosts is essential. This study presents a method for predicting potential hosts for plasmids and offers insights into the potential paths for spreading functional genes between different bacteria. Understanding plasmid-host relationships is crucial for comprehending the ecological and clinical impact of plasmids and implications for various biological processes.

## INTRODUCTION

Plasmids are extrachromosomal, mainly circular, DNA elements, common in Bacteria, Archaea, and some Eukarya (1). They are important biological elements at multiple levels: they are carriers and spreaders of antibiotic resistance (2–(4); they facilitate the environmental adaptation of bacteria to new stresses (5–(7); they play a significant role in bacterial evolution (8–(10); and they also serve as valuable biotechnological tools, being used as cloning and expression vectors (11, 12). However, plasmids also impose a burden on the host and, therefore, come with a fitness cost (13). This cost leads to the evolution of various bacterial defense systems against plasmids, such as the CRISPR (Clustered Regularly Interspaced Short Palindromic Repeats) system (14) and the Wadjet system (15).

Traditionally, plasmids were identified from isolates or isolated via conjugation to a culturable bacteria. While these methods provide some information about the natural host (in the case of isolate) or a potential host (in the case of conjugation), there is a need for further studies to achieve information about other potential hosts and host range (16). Lately, the availability of high-throughput sequencing technologies at relatively low prices for environmental metagenomics studies is generating increasing numbers of plasmidome studies covering a variety of environments (17–(23). However, irrespective of the data collection and sequencing methods, typical metagenomic data fails to associate plasmids with their bacterial hosts, because the data do not contain direct information about the cellular context of the plasmidic extrachromosomal DNA sequences (16).

Knowing the bacterial host of a plasmid is potentially important information in view of the crucial role played by plasmids in horizontal gene transfer (HGT) and in spreading functional genes among bacterial populations. Similarly, there is also a need for information on the host range of plasmids, especially when attempting to understand the effects of plasmids on a bacterial population (24–(26). In addressing this need, Redondo-Salvo et al. introduced a scale for ranking the host range of plasmids, between Grade I for plasmids that can reside in species within a single genus to Grade VI for plasmids found in different phyla (27). In their analysis of common plasmid taxonomic units (PTUs), Redondo-Salvo et al. found that mobilizable and conjugative plasmids (designated MOB+ plasmids) exhibited broader host distributions than non-mobilizable plasmids (designated MOB-plasmids). However, conjugation is not the only mechanism for plasmid dissemination, and other mechanisms, such as transformation (28, 29) and transduction (30), were also shown to enable plasmid transfer between bacteria, even between bacteria of different taxonomic classifications.

In recent years, there have thus been several attempts to experimentally collect data that contain information about both the plasmids and the host. One approach is based on Hi-C methods (16, 31–33), in which adjacent DNA molecules are crosslinked before cell lysis and DNA fragmentation, thereby keeping together DNA fragments that are in close proximity in the cell (such as chromosomal DNA and plasmids, at least in some cases). Using a different approach, Beaulaurier et al. used DNA methylation profiling to identify similar methylation patterns that could assign plasmid DNA to a host (34). Recently, progress in the development of single cell sequencing methods (35) has enabled the identification of plasmids in the context of the bacterial host. However, all these methods are expensive, require specialized skills and equipment, and exhibit low sensitivity and low throughput. A more affordable and scalable approach would be to associate plasmids with their potential hosts *in silico*, as commonly done when studying viruses and their hosts.

A parallel research direction in bacteriophages and bacterial viromes also addresses host prediction, and several methods have been developed for the prediction of hosts for bacteriophages. As plasmids and viruses share some basic features in their interaction with the bacterial host, concepts that have been applied for host prediction in viruses may also be useful (with appropriate adaptation) for plasmids. Efficient approaches to detecting hosts of viruses include both methods based on sequence signatures, typically by calculating similarities between the phage sequence and each potential host genome by oligonucleotide frequency, Markov chain model or Gaussian mode (36–(42), and alignment-based methods (43). These methods provide host prediction from standard metagenomic data, without a need for additional experimental protocols.

In fact, the current *in silico* methods for plasmid hosts are based on concepts similar to those used for predicting virus hosts. Methods which are based on phylogenetic association (44) or genomic signatures analysis that compares the distance in the dinucleotide composition between the plasmid and chromosome sequences (45, 46) were reported. Since these approaches are based on genetic exchange and coevolution of the plasmids with the host chromosome, they provide good host prediction for narrow-host plasmids and long-term hosts, but they are limited for broad-host plasmids. Recently, Aytan-Aktug et al. developed a machine-learning-based tool, the PlasmidHostFinder, that uses a set of random-forest-based models for predicting plasmid hosts at different bacterial taxonomic levels (47). PlasmidHostFinder, which uses pattern detection methods rather than homology-based methods, provided higher accuracy in host prediction, although sensitivity was still low.

The current study utilizes the bacterial CRISPR system for the prediction of potential plasmid hosts. Bacterial CRISPR is a well-recognized bacterial adaptive immune system against foreign DNA molecule, such as plasmids, viruses, and other mobile genetic elements (42, 48, 49). While there are various CRIPR-Cas systems which are classified into two major classes, six types and more than 45 subtypes (50), all CRIPR-Cas systems share the same fundamental mechanism of cutting foreign DNA into short fragments (26 to 72 bps) and integrating them as spacers between repeat sequences to create a memory for future encounters [for further details about CRISPR-Cas systems, please refer to (51–(53)]. Hence, the CRISPR spacers act as a ‘record’ for historic encounters between viruses or plasmids and their potential hosts (54–(57). As such, CRISPR spacers have been shown to be effective as predictors of potential hosts for viruses (58–(60). Recently CRISPR spacers was also used as predictor of hosts for plasmids (61). However, unlike the case of viruses, the use of CRISPR spacers as host predictors for plasmids has not been systematically studied yet.

Plasmids are known for their dynamic nature and have a propensity to exchange genetic material, particularly accessory genes, with various genetic elements, including chromosomes, phages, or other plasmids (62, 63). Yet, it remains uncertain whether a spacer that matches a plasmid was originally acquired from plasmid specific regions, making the use of the spacers as host predictors feasible. Otherwise, in case most spacers that match plasmids are acquired form exchangeable accessory genes, CRISPR spacers may not serve as good host predictors for plasmids. Nonetheless, previous studies have demonstrated that, in the case of viral infections, spacers are acquired from early injected genomic regions (64). As a result, the targets of the CRISPR-Cas system are not randomly dispersed across the viral genome but are notably concentrated in locations upstream of the *cos* site (64). While it is currently unknown whether a similar preference for specific genomic regions exists in plasmids, a comparable bias in plasmids could potentially impact the ability to predict the host based on spacers.

In leveraging the CRISPR-Cas system for plasmid host prediction, one should be aware of the drawback that the predictions would be fundamentally limited to hosts that contain CRISPR-Cas defense systems in their genomes. Indeed, based on an analysis of bacterial and archaeal genomes in currently available databases, Makarova et al. reported that CRISPR-Cas is found in only 40% of bacteria and 81% of archaea (50). These numbers agree with previous studies (65–(67) and are commonly cited as the actual prevalence of CRISPR-Cas in microorganisms. Nevertheless, by analysis of a large environmental data set, using a cultivation-independent approach, Burstein et al. estimated that only 10% of microorganisms contain a CRISPR-Cas system and showed that some major bacterial lineages do not contain CRISPR-Cas defense system at all (68). The above notwithstanding, our working hypothesis for this study was that a CRISPR-Cas-based methodology could be applied, at least partially for the microorganisms which contain CRISPR-Cas system, for predicting potential hosts for plasmids.

Here, we test the use of the alignment of CRISPR spacers to predict potential bacterial hosts for plasmids. We show that the CRISPR-based method is efficient and accurate. We predict potential hosts for 46% of the plasmids in the Plasmid Database (PLSDB) (69) and captured the reported host in 84% of the cases at the family level and 99% of the cases at the phylum level. Moreover, our findings indicate that CRISPR spacers are predominantly acquired from plasmid-specific backbone genes, while functional genes, that may be shared with other genetic elements, remain untargeted. Importantly, this method is not restricted to narrow-host plasmids and can also predict the host range of plasmids.

## RESULTS

### Testing the hypothesis

Our hypothesis is that by aligning a plasmid of unknown host to the spacers in the CRISPRCasdb, we can predict potential hosts for the unknown plasmids. In order to test it, we aligned all the plasmids in the PLSDB (69) to all the spacers from the CRISPRCasdb (70) (Fig 1a). The alignment yielded 730,841 hits of 15,891 out of 34,513 (46%) plasmids (“matched plasmids” hereafter) to 3,181 different bacterial strains of potential hosts. The average number of hits per plasmid is 46, with a standard deviation of 115. The 25th, 50th, and 75th percentiles are 2, 5, and 25, respectively (full distribution see Fig. S1). Such a skewed distribution is an indication of the presence of a few plasmids with a high number of hits. The maximum hits per plasmid is 1,403 hits on plasmid accession number NZ_LR134400, a 593,685-bps plasmid isolated from Listeria monocytogenes. However, 1,385 of these 1,403 hits are from different isolates of Listeria monocytogenes, representing hits against different targets on the plasmid and hits to the same target from different isolates of Listeria monocytogenes. The remaining hits are from Lactobacillus acidophilus, which also belongs to the Bacilli class, and from Klebsiella pneumoniae and Neisseria lactamica, which are Proteobacteria. Hence, this plasmid is a broad-host plasmid that can colonize hosts from different phyla.

**Fig. 1.**
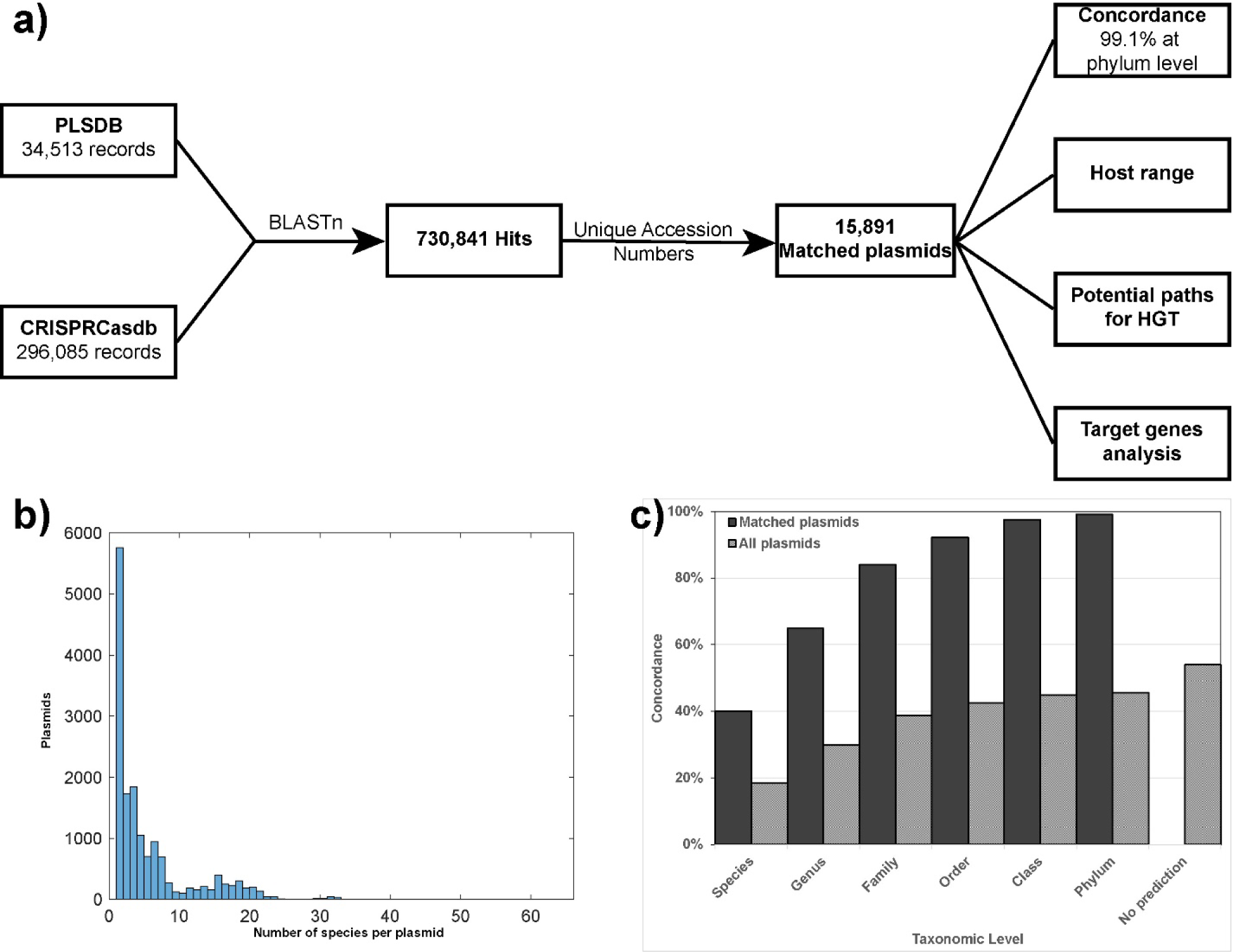
The CRISPR-based method and its results. a) Schematic representation of the basic method and the additional analysis and information that can be extracted. b) The distribution of number of species per plasmid. c) Concordance of the predictions of the CRISPR-based method with the reported host at each taxonomic level; for each taxonomic level, the percentage of matched plasmids (solid columns) and of all the plasmids in the PLSDB (dotted columns) for which the prediction of our method included the known host in the database is shown.

The next two plasmids with a high number of hits are NZ_CP079158.1 and NZ_CP079675.1, which are practically the same plasmid (99% coverage and 99.99% identity) isolated from Klebsiella pneumoniae, both having 1,156 hits. All the hits represent different targets on the plasmid and different isolates of Klebsiella pneumoniae or the closely related Klebsiella variicola, therefore, despite the large number of hits, they correspond to a narrow host range plasmid.

To get a better understanding of the specificity of the hosts prediction, we looked at the number of predicted host species per plasmid (Fig 1b), rather than hits per plasmid mentioned above. For all matched plasmids, we found that on average, each plasmid is found in 5.19 species, with a standard deviation of 5.96, and 25th, 50th, and 75th percentiles of 1, 3, and 6, respectively. The maximum number of species per plasmid is 63, in the case of plasmid CP053320.1. However, when manually checking the predicted hosts, we found that all the 63 different predicted host species of CP053320.1 are subspecies of Salmonella enterica. Moreover, plasmid NZ_LR134400, which is mentioned above as the plasmid with the highest number of hits (1,403), is predicted to be associated with only 16 different species, but 11 of them are subspecies of Listeria monocytogenes.

Overall, the results shown above indicate that while we allow the prediction of multiple potential hosts with the aim to increase the knowledge about alternative hosts and host range, the results are mostly confined to a limited number of hosts, suggesting that the targeted sequences are not random sequences which are common to large number of plasmids.

### Concordance with the known host

To evaluate the ability of the CRISPR-based method to capture the correct host for the matched plasmids, the concordance of the method was estimated by comparing the predictions of the method with the known hosts of the matched plasmids. For each matched plasmid, the hits that yielded the best match to the known host, namely, the lowest taxonomic level at which the prediction agrees with the known host, were identified (Fig. 1c). The reported host at the species level was included in our predictions for 6,363 plasmids (40.0% of matched plasmids, 18.4% of all plasmids in the database). For an additional 3,937 plasmids, the reported host was included in our prediction at the genus level (total of 10,300, 64.8% of matched plasmids, 29.8% of all plasmids). As expected, the concordance increased as the taxonomic level went up, with 13,356 of the reported hosts included in our predictions at the family level (84.0% of matched plasmids, 38.7% of all plasmids), 14,658 of the reported hosts included in our predictions at the order level (92.2% of matched plasmids, 42.5% of all plasmids), 15,493 of the reported hosts included in our predictions at the class level (97.5% of matched plasmids, 44.9% of all plasmids), and 15,751 of the reported hosts included in our predictions at the phylum level (99.1% for matched plasmids, 45.6% of all plasmids), as shown in Fig. 1c. Only 140 predictions (0.9% of matched plasmids) did not include the predicted host, even at the phylum level. However, 60 of these predictions (42.9%) are classified as ‘uncultured bacterium’ in the PLSDB, and, therefore, they may be considered as predicted hosts rather than mismatches (for example, see the case of MG879028.1 discussed below). The method could not predict a host for 54.0% of the plasmids in the PLSDB, but this outcome was expected considering the limitations of the method (see Discussion).

### Host family distribution

Most of the hits (681,995 of 730,841; 93.3% of hits) were obtained for the Enterobacteriaceae family. The second and third families with high numbers of hits were Xanthomonadaceae (9,112 of 730,841; 1.2% of the hits) and Lactobacillaceae (4,106 of 730,841; 0.56% of the hits), respectively. At the class level, 96.4% of the hits (704,873 hits) were obtained for Gammaproteobacteria, which is the most studied class, containing some of the best studied pathogens and laboratory strains having many sequenced isolates. For that reason, the number of hits from Gammaproteobacteria is inflated by isolates matching the same plasmids.

A better evaluation of the distribution at the family level would be the evaluation of the distribution of the taxonomic families of the reported host for the matched plasmids. That way, we overcome the inflation in the number of hits due to multiple sequenced isolates of the same species, although we still suffer from the bias of the databases. Also here, Enterobacteriaceae is the most abundant family, corresponding to 10,858 of the 15,891 matched plasmids (68.3%), but the other abundant families are Staphylococcaceae (930 of 15,891, 5.9%), Lactobacillaceae (834 of 15,891, 5.2%) and Enterococcaceae (771 of 15,891, 4.9%), as shown in Table 1. At the class level, 75.0% (11,913) of the reported hosts of the matched plasmids are Gammaproteobacteria, and 19.1% (3,034) were found in Bacilli. When examining only plasmids for which the method included the reported host at the species level (the most successful cases), the dominance of the Enterobacteriaceae family was even more pronounced, standing at 83% (5,279 of 6,363; Table 1). The high percentage of specific families in both the total hits and the matched plasmid groupings is an indication of the bias of our method and the databases toward the more studied microorganisms and toward CRISPR-Cas containing families.

**Table 1.** Families of the reported hosts of the matched plasmids.

| All matched plasmids |  |  | Plasmids with accurate prediction at the species level |  |  |
| --- | --- | --- | --- | --- | --- |
| Family | No. of plasmids | Percentage | Family | No. of plasmids | Percentage |
| Enterobacteriaceae | 10858 | 68.33 | Enterobacteriaceae | 5279 | 82.96 |
| Staphylococcaceae | 930 | 5.85 | Enterococcaceae | 250 | 3.93 |
| Lactobacillaceae | 834 | 5.25 | Lactobacillaceae | 148 | 2.33 |
| Enterococcaceae | 771 | 4.85 | Moraxellaceae | 68 | 1.07 |
| Streptococcaceae | 258 | 1.62 | Staphylococcaceae | 54 | 0.85 |
| Xanthomonadaceae | 216 | 1.36 | Pseudomonadaceae | 52 | 0.82 |
| Moraxellaceae | 190 | 1.20 | Bacillaceae | 44 | 0.69 |
| Bacillaceae | 188 | 1.18 | Streptococcaceae | 43 | 0.67 |
| Yersiniaceae | 169 | 1.06 | Peptostreptococcaceae | 40 | 0.63 |
| Burkholderiaceae | 133 | 0.84 | Campylobacteraceae | 38 | 0.60 |
| Pseudomonadaceae | 97 | 0.61 | Clostridiaceae | 30 | 0.47 |
| Other | 1247 | 7.85 | Other | 317 | 4.98 |
| **Total** | **15891** | **100** | **Total** | **6363** | **100** |

### Host-range prediction

Importantly, 77.5% (12,316) of the matched plasmids had more than a single hit, suggesting that the CRISPR-based method can indeed be used for predicting the host range of the plasmids. The host range scale introduced by Redondo-Salvo et al. (27) was used to assign each matched plasmid to a host range grade based on the predictions of CRISPR-based method. Of the matched plasmids, 59.4% (9,433 plasmids) were found to belong to Grade I (Fig. 2a, left bar), i.e., they may vary only at the species level. Of these Grade I plasmids, 62.1% (5,858) matched only to a single species, with 3,575 of them having only one hit. The percentages of plasmids assigned to the other host range grades were 25.8%, 5.8%, 6.9%, 0.8% and 1.5% for Grades II, III, IV, V and VI, respectively. The average grade for all the matched plasmids was 1.68, as most of the plasmids are narrow range and belong to Grades I and II.

**Fig. 2.**
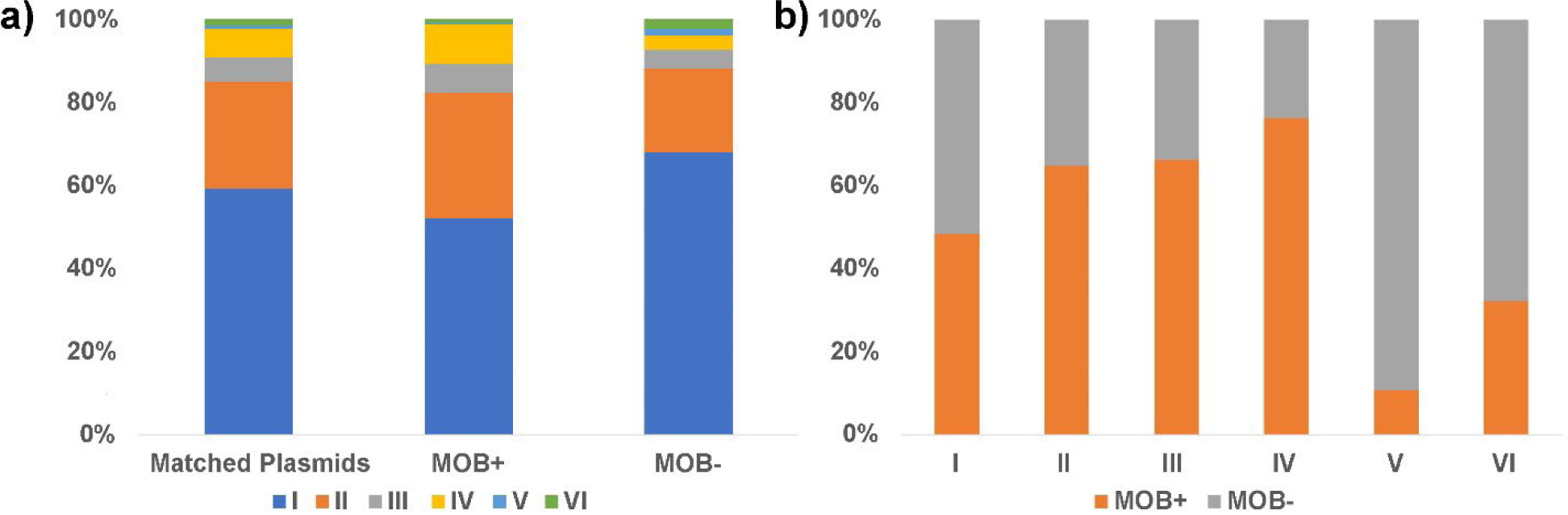
Host range grades for mobile and non-mobile plasmids. a) Percentage of each host range grade in all the matched plasmids (all, left bar), in mobile plasmids (MOB+, middle bar) and in non-mobile plasmids (MOB-, right bar). b) Percentage of mobile (MOB+, orange) and non-mobile (MOB-, gray) plasmids in each host range grade.

Separating the host range of the MOB+ plasmids from that of the MOB-plasmids showed that, on average, MOB+ plasmids tend to be broader in their host range compared to MOB-plasmids, especially at the family level (Grade III) and the order level (Grade IV), with an average grade of 1.78 for MOB+ versus 1.57 for MOB-(Fig. 2a, middle and right bars; Fig. 2b). However, surprisingly, at the class level (Grade V) and phylum level (Grade VI), the trend was reversed, and MOB-plasmids showed broader distribution across the different classes (p value <0.001) and phyla (p value <0.001) (Fig. 2b).

### Network analysis of matched plasmids

The output of the CRISPR-based method can also be presented as a network of the connections between plasmids and host families (Fig. 3). This network shows a potential flux of genetic material between different taxonomic levels via plasmids. As expected, most of the connections are within-phylum, with only a few plasmids connected to hosts from more than one phylum, thus allowing cross-phylum transmission of genetic material. Since our study is based on known plasmids in the database, it is strongly biased toward Proteobacteria and Firmicutes, which are the most studied phyla and thus the most represented in the CRISPR database. Plasmids connected to these two phyla mostly share hosts between families within the same phylum (large ellipses in Fig. 3), with some cross-phylum interactions. However, four families in the Firmicutes phylum appear within the region of the Proteobacteria (green rectangles in the red ellipse). In the case of two of them, Listeriaceae and Oscillospiraceae, the number of connected plasmids is relatively low (11 and 2, respectively) and since one of these plasmids is a broad host plasmid, the families appear to share plasmids with the Proteobacteria phylum. In contrast, the two other Firmicutes families in the Proteobacteria ellipse, Bacillaceae and Paenibacillaceae (arrows in Fig. 3), have high number of plasmid connections (166 and 60, respectively) and yet they are relatively isolated from the other the Firmicutes families. The Bacillaceae family is relatively isolated, with only one cross-family connection, which is also a cross-phylum connection, to Mesomycoplasma of the Tenericutes phylum via plasmid pDSM15939_1 (accession number NZ_CP015439.1) isolated from *Anoxybacillus amylolyticus* strain DSM 15939 (71). Nevertheless, the Paenibacillaceae family is highly connected via multiple plasmids to families of the phyla of Proteobacteria (Gamma and Beta) and Tenericutes, but not connected to any other Firmicutes families.

**Fig. 3.**
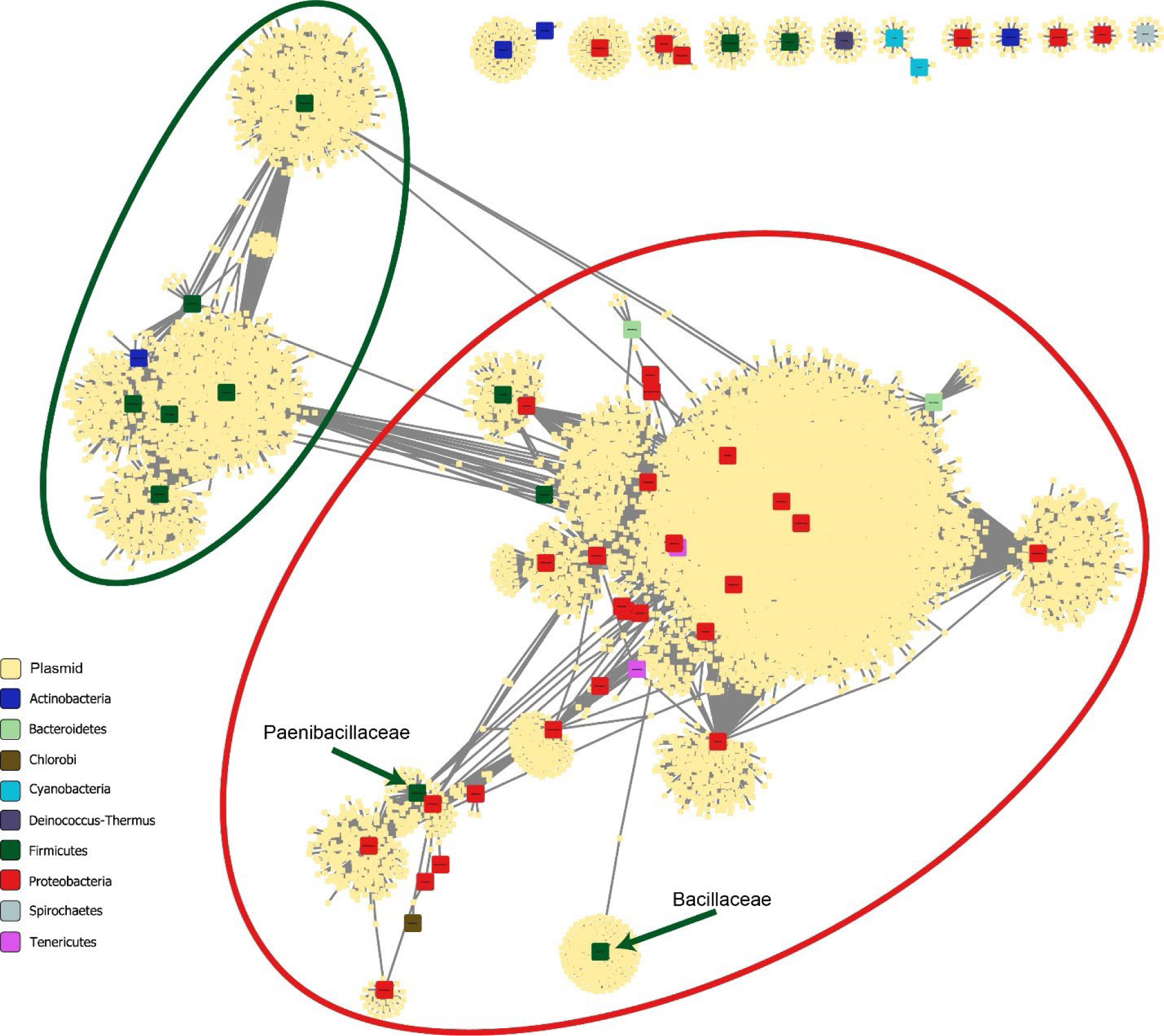
Connections between different bacterial families via plasmids. Each large colored square represents a bacterial taxonomic family, small yellow squares represent plasmids, gray lines connect between plasmids and the families of their predicted hosts. The families are colored according to the phylum. The large ellipses represent the region of the Proteobacteria (red) and Firmicutes (green). Families that have less than 10 plasmids and are not connected (via plasmids) to another family were eliminated from the diagram.

### A case study: MG879028.1

An example of the predictive capabilities of the method is the plasmid pEG1-1 (accession number MG879028.1), which was isolated from uncultured bacterium from a sample of environmental sediments (72). The pEG1-1 plasmid contains genes for heavy metal resistance, agricultural antibiotic resistance, and human antibiotic resistance. It was assumed to be a broad host plasmid, as, according to its sequence analysis, it belongs to the IncP-1β group, which is known to be a broad host group (72). It is a conjugative plasmid, containing two conjugation modules. It also contains the in104 complex integron, which has been demonstrated only in a few genera of the Enterobacteriaceae family, and therefore Enterobacteriaceae was assumed to be its natural host. In our study, however, hosts of the Enterobacteriaceae family were not predicted, but hosts from other Gammaproteobacteria families were predicted - Pasteurellaceae, Pseudomonadaceae and Xanthomonadaceae, as well as the family of Comamonadaceae (Betaproteobacteria class) and Paenibacillaceae (Firmicutes phylum), thereby confirming its broad host nature (host range Grade VI).

### PTU network

PLSDB is based on deposited sequences from various isolates and studies. However, it is possible that several records in the PLSDB refer to different versions of the same plasmid. Therefore, whenever applicable, we assigned PTUs to the plasmids and redid the network analysis at the PTU level. As only a small fraction of the plasmids was assigned to PTUs (2,722 plasmids out of the 15,891 matched plasmids), the network at the PTU level was not as rich as the network at the plasmid level, but the trend was similar, namely, the separation between families of the Firmicutes phylum and families of the Proteobacteria phylum was still observed, with the Bacillaceae and Paenibacillaceae families standing out as being disconnected from the other families of the Firmicutes phylum (Fig. 4). As discussed above for the plasmid level, at the PTU level the Bacillaceae family is also isolated from other families and does not share plasmids with other families, while the Paenibacillaceae family does share plasmids with other families of the Proteobacteria phylum via the broad host PTU-P1. PTU-P1 was described by Redondo-Salvo et al. as the broadest host PTU, with plasmids from this PTU being found scattered throughout the entire bacterial kingdom (27).

**Fig. 4.**
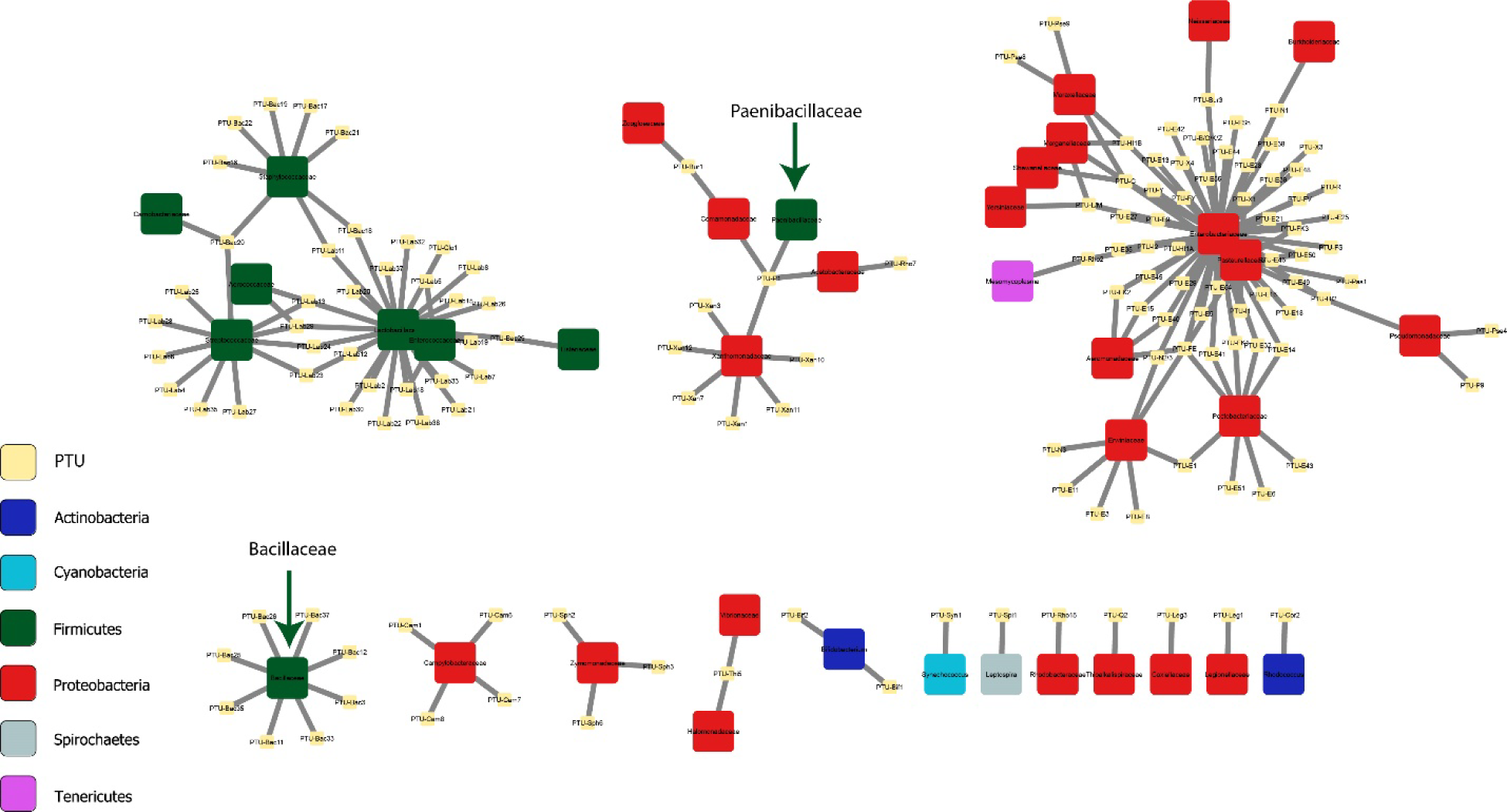
Connections between different families via PTUs. Large colored squares represent taxonomic families and are colored according to the phylum. Small yellow squares represent PTUs. The edges represent the presence of the PTU in the connected family.

### Spacers bias toward specific genetic regions on the plasmids

A match of a CRISPR spacer to a plasmid can occur due to a specific spacer against the plasmid or due to a spacer against an insertion sequence, which may originate from a different source and not necessarily represents a previous encounter of the predicted host with the plasmid. The latter case may produce artifacts regarding the predicted hosts and host range. Therefore, we analyzed the annotation as it appears in the NCBI database in an attempt to recognize insertion elements. Only 364,400 of the 730,841 hits (49.9%) fall within an annotated feature. Among these, only 25 hits, coming from 9 plasmids, are annotated as falling within a feature type of ‘mobile_element’, namely, annotated as located within an IS element. Moreover, all the 9 plasmids that correspond to the 25 hits are ranked by our analysis as host range grade I or II and show concordance with the reported host at the species, genus, or family taxonomic level. This means that the IS did not result in a higher number of hits from hosts all over the taxonomic scale.

However, the annotation of mobile elements and IS elements in the NCBI database is incomplete, necessitating further analysis of the target gene annotations. Consequently, we calculated the frequencies of all target gene annotations and manually categorized these genes based on their likelihood of being plasmid backbone genes or IS elements (see Fig. 5, for complete list of target genes see Table S1). Approximately one-third of the annotated genes (31.8%) are labeled as ‘hypothetical protein’ or ‘DUF’ (Domain of Unknown Function). However, upon closer examination of the annotated target genes and domains with known functions, we determined that 45.2% of the plasmid-derived spacers aligned with genes that unequivocally qualify as plasmid backbone genes. These include genes specific to plasmid backbones, such as conjugation genes, plasmid replication control genes, and plasmid-specific addiction systems. An additional 43.1% ORFs are genes with typical plasmid functions, albeit not exclusive to plasmids, primarily involving general DNA modification and processing functions. Only 2.9% of the annotated target genes are IS genes, such as transposases and integrases, while 8.8% are general functional genes with functions not specifically associated with plasmids. These can be either functional genes within the plasmid’s backbone or genes originating from IS elements. The predominance of plasmid-related annotations among the annotated genes serves as additional evidence that the matches between CRISPR spacers and plasmids identified in this study likely result from previous interactions between the predicted host and the plasmid. These findings argue against artifacts arising from mobile elements, which may have led to spacer production against sources other than the plasmid. Interestingly, none of the annotated target genes is a gene related to antibiotic or metal resistance.

**Fig. 5.**
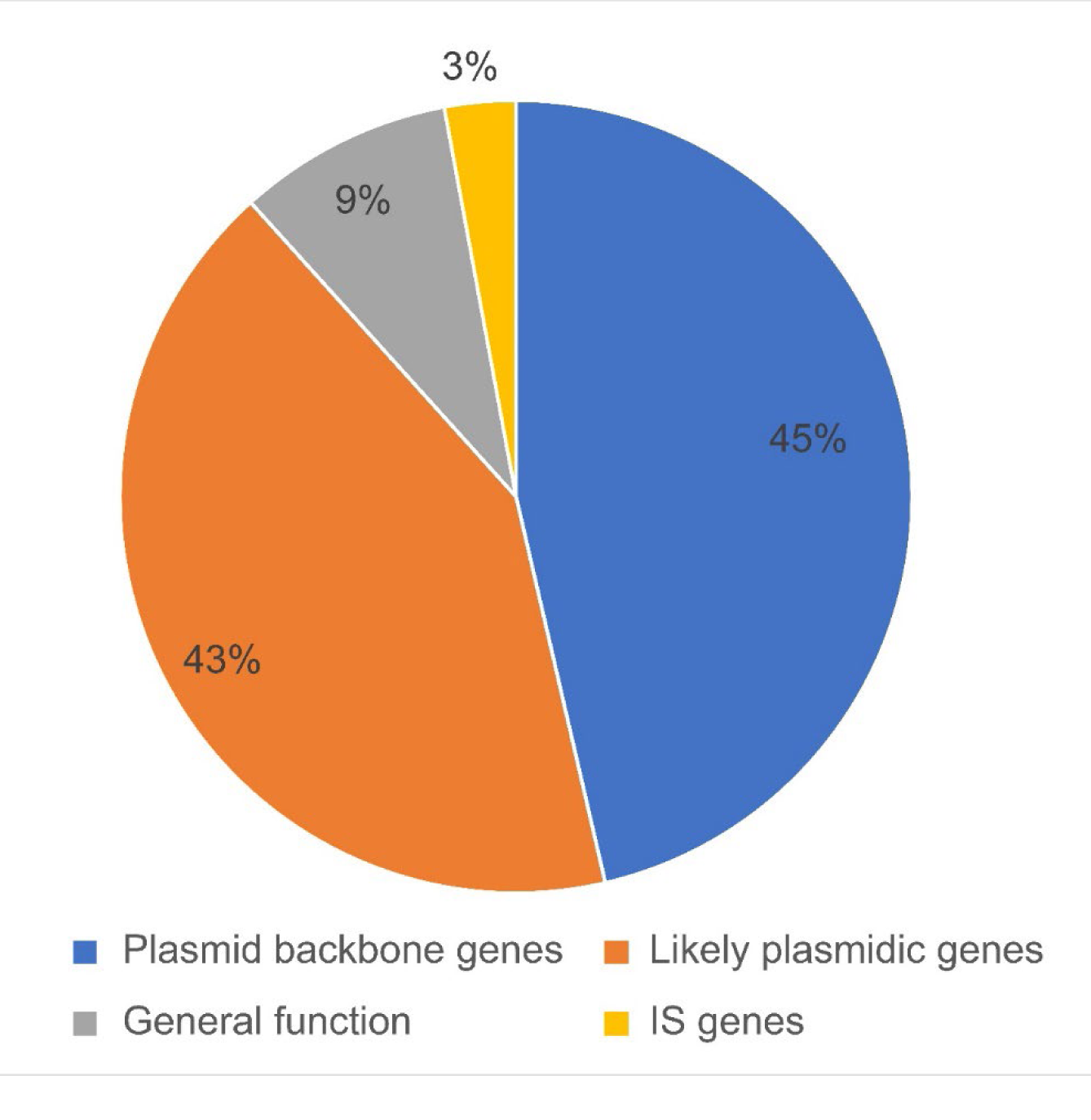
Analysis of the targets on the plasmids of CRISPR spacers. Targets that have annotation for a known function were classified into four groups: genes which are clearly plasmid backbone genes (blue), genes which have a function that is typical for plasmids but is not specific (orange), genes with general function (grey) and genes with IS related function (yellow).

## DISCUSSION

The ability to predict hosts for viruses using CRISPR spacers has been demonstrated and tested with various available tools based on this approach (58–(60). Recently, several studies have employed the alignment of plasmid sequences to CRISPR spacers to predict hosts for the plasmids (61). However, it is not clear whether this method would be as efficient and accurate for plasmids as it is for viruses. Plasmids are known for their dynamic nature, frequently exchanging genetic material with other genetic elements within the host cell (62, 63). Therefore, it is possible that a spacer acquired from one genetic element could target a plasmid containing the same sequence, even if the host cell has never encountered the plasmid. If this is the case, CRISPR spacers are expected to be inefficient and inaccurate.

In this study, we tested the use of CRISPR spacers as potential predictors for plasmid hosts by comparing the reported host(s) for the plasmids in the PLSDB with the predicted host(s) from the CRISPR-based method. We also demonstrated the utility of this method in extracting additional information, such as host range and the potential network of gene transfer. This approach is fundamentally limited to the identification of bacterial hosts that accommodate CRISPR-Cas defense systems in their genome. Taking the fundamental limit of such method into account, the ability of our method to predict bacterial hosts for 46.0% of plasmids from the PLSDB in concordance with the reported host of 84.0% of hosts at the family level and more than 99% of hosts at the phylum level (Fig. 1c) is encouraging and indicates the usefulness of the method. Overall, for 45.6% of the plasmids in the PLSDB the reported host at the phylum level was included in our predictions. Our analysis suggests that the high concordance is not due to a large number of predicted hosts per plasmid. By comparison, the phylogenetic association method that was developed by Kav et al. achieved only 27.8% accuracy at the phylum level (44), although that method is not limited to CRISPR-Cas-containing microorganisms.

A possible explanation for our relatively high success rate may relate to the fact that both PLSDB and CRISPRCasdb are highly biased and enriched by plasmids and spacers from well-characterized bacteria. This bias is also reflected in the dominance of the order Gammaproteobacteria and, more specifically, the family Enterobacteriaceae, in the predictions. Burstein et al. stated that the NCBI database is biased toward CRISPR-Cas-containing bacteria and showed that entire lineages of uncultivated organisms, which are underrepresented in the database, are essentially devoid of CRISPR-Cas (68). Therefore, our test dataset is enriched with plasmids from hosts that most probably possess CRISPR-Cas systems, although it is still limited to the reported prevalence of 40% of bacteria and 81% of archaea (50). It will be informative to revisit this analysis as the database becomes enriched with more uncultivated organisms to examine whether a larger dataset will increase the strength of the method by virtue of a larger repertoire of CRISPR spacers or whether it will reduce the hit rate due to the larger percentage of plasmids isolated from microorganisms that are devoid of CRISPR-Cas systems.

Plasmids have a dynamic nature, and they often exchange genes with their bacterial hosts’ chromosomes, viruses, and other plasmids (62, 63). This exchange may impose another limit on the CRISPR-based method, as some CRISPR spacers may be directed against a sequence from one genetic element (plasmid or virus) but also match an identical sequence on a different plasmid. This would lead to a match between the latter plasmid and the host of the former genetic element, even if that host had never encountered that plasmid. However, the analysis of the CRISPR targets suggests that, in most cases, the target is a plasmid-specific target rather than an IS sequence that may have originated from a different mobile element. Strikingly, we did not find any spacer targeting antibiotic resistance or metal resistance genes. Such avoidance can be a result of a negative selection against resistance genes as spacers, or a result of the biological mechanism of spacers acquisition, as shown in the case of viruses (64). Regardless of the origin of this bias, the fact that most spacers target plasmid specific genes increases our confidence in the hosts predictions obtained by the method.

Moreover, the ability of our method to capture the correct host at the phylum level for more than 99% of the plasmids that had matches in the spacers database is another indication that the predictions are not due to general genetic elements which are shared by many plasmids (and viruses and chromosomes) but rather more specific spacers for the targeted plasmid. The relatively low average and median of predicted species as potential hosts per plasmid (5.19 and 3, respectively) is another indication that the predicted hosts and host range are not the result of spacers targeting IS elements that can be found all over the bacterial range, but rather the result of more specific targeting by the CRISPR system. Nonetheless, similar to any other computational prediction, the host predictions generated by this method necessitate further validation when investigating specific plasmids.

One could improve the CRISPR-based method by introducing a score for each predicted host, based on the nature of the target sequence, the nature of the target region and the number of spacers from the host against the plasmid. Moreover, additional selection steps could be added to increase sensitivity and accuracy, such as comparing spacers and plasmids at the protein level and combining evidence from multiple spacers matching the same plasmid (59), or adding a filtering step inspired by the biologically relevant function of the CRISPR-Cas systems (58). On top of that, it might be possible to enhance the CRISPR-based method by integrating it with other approaches for identifying plasmid hosts, such as a method based on phylogenetic association (44) or genomic signatures (45, 46). For environmental datasets collected from different niches, one may refine the host prediction by comparing with colocalization data of plasmids and bacteria. Although these additional steps may improve the method’s accuracy and enable the selection of the most probable host, the CRISPR-based approach by itself, despite its simplicity, seems to effectively fulfill its purpose by capturing the reported host. Furthermore, the method doesn’t restrict predictions to the most likely host but includes all potential hosts, thereby predicting alternative potential hosts and estimating the host range.

Interestingly, 3,387 out of the 6,363 (53.2%) matched plasmids which are in concordance with the reported host at the species level were also assigned to other species, suggesting potential alternative host species. Similarly, the assignment of the other 9,528 plasmids to a species different from the species from which they were isolated does not necessarily indicate a failure of the method, but rather a prediction of a potential alternative host. The finding that most of these “misassigned” plasmids were assigned to species within the same family as the reported host strengthens the possibility that these “misassignments” are, in fact, potential alternative hosts. While these predictions should be taken with caution, as described above, they indicate that the method is not only useful for predicting the preferred and/or the most common host but also allows the prediction of the host range of the plasmids.

It is commonly accepted that mobile and conjugative plasmids tend to have broader host range than the non-mobile plasmids (27). Indeed, our results indicate that, in general, mobile plasmids present higher average host range grade (1.78 for MOB+ vs. 1.57 for MOB-). However, at host range Grades V and VI, which are the broadest categories, we found more non-mobile plasmids. While this may seem surprising at first, a possible explanation is the presence of alternative transfer mechanisms, such as transformation (28, 29) and transduction (30), in addition to conjugation.

These alternative mechanisms are more “passive” than conjugation, and hence do not require a specific interaction between the transfer machinery and the new host. For that reason, passive mechanisms may allow the introduction of plasmids into bacteria that are further away in the phylogenetic tree. If this is the case, it might be speculated that the prevalence of transfer mechanisms other than conjugation is higher than commonly thought.

The plasmid-host network shows that, as expected, most of the plasmids colonize hosts from a single bacterial family, and most of the cross-family interactions are within the same phylum. Yet, some plasmids colonize hosts from different phyla, suggesting a potential cross-phylum genetic transmission. This mode of genetic transmission is clearly visible within the families of the Firmicutes and Proteobacteria phyla, which are the best studied phyla and therefore represent most of the hosts found in our study. These two phyla produced two separate regions in the connectivity network with very little cross interaction. Yet, the Bacillaceae and Paenibacillaceae families, which belong to the Firmicutes phylum, do not share plasmids with other families in their phylum. Bacillaceae seems to be isolated from all other families, while Paenibacillaceae is connected to various families, but all from different phyla. The reason for the uniqueness of these families is not clear to us at this point and further study is needed to verify and understand it.

The case of plasmid pEG1-1 with accession number MG879028.1 is presented as an example of the use of our method. This plasmid was isolated from an unculturable bacteria from an environmental sample. The CRISPR-based method provides predictions for its potential hosts and its host range. While our findings are still predictions until validated, they are in agreement with the knowledge we have about this plasmid as being a broad host IncP-1β group plasmid, which is expected to be found in hosts belonging to the Gammaproteobacteria order.

## CONCLUSIONS

In summary, we have shown that CRISPR spacers can be used for the prediction of potential hosts for plasmids efficiently. It is shown to effectively capture the reported host at the family level (84%) or the phylum level (99%), among additional potential hosts in some cases, and it can also be used to predict alternative hosts and the host range for known plasmids. Notably, CRISPR spacers primarily target plasmid backbone genes, as opposed to the more dynamic functional genes. This selective targeting pattern of spacers boosts our confidence that the spacer was acquired through a prior encounter with the plasmid. Using our predictions, we compared the host range of mobile plasmids with that of non-mobile plasmids. While mobile plasmids are less species specific and tend to have broader host range up to the order level, at the class and phylum levels the non-mobile plasmids are more diverse, suggesting the higher-than-expected contribution of transfer mechanisms other than conjugation in HGT across non-related bacteria. Nevertheless, one should keep in mind the limitations and the bias of the method, and therefore conclusions should be taken with caution before further validation. Our results clearly show that despite the limitations (can only predict hosts that use CRISPR-cas systems and use database with a strong bias toward specific taxa), the CRISPR-based method is able to accurately predict the host of almost half of the known plasmids.

## METHODS

### Datasets

All 34,513 records from the PLSDB (69) v. 2021_06_23 and all 296,085 spacers from CRISPRCasdb (70) version 20210121 were downloaded and used as is, without additional filtration or curation. All codes were written in Python version 3.8 unless otherwise noted and are available on https://github.com/Tal-Lab/crispr_plasmidome.

### Alignment of plasmids to spacers

We utilized the Bio.Blast.Applications module to convert the spacers fasta file from CRISPRCasdb into a spacers database using the makeblastdb command. Then, we employed BLASTn to align the plasmids from the PLSDB to the spacers database (100% identity, E-value 10^-5^, no gaps). To determine the appropriate percent identity thresholds, we conducted tests at 90%, 95%, and 100% cutoffs. Surprisingly, no significant differences were observed between the various cutoffs (refer to Supplementary Fig. S2 and S3 for results when using a threshold of 90% identity). Given the lack of significant advantages in reducing the threshold, we opted to use a threshold of 100% identity for this manuscript. It is important to note that plasmids can have multiple hosts, and since the method also serves host range prediction, we retained all the predicted hosts without selecting the most probable one.

In cases where a plasmid was aligned to a spacer from the same plasmid (i.e., same accession number for the plasmid from the PLSDB and the host of the spacer), we removed that record, as it indicates a plasmid-borne spacer rather than a spacer targeting the plasmid.

### Finding the concordance level

For each spacer-plasmid hit, the full taxonomic lineage of the plasmid was extracted from the PLSDB. The strain of the matching spacer was obtained from the CRISPRCasdb, and the full taxonomic lineage of the potential host was retrieved using the Entrez Programming Utilities (E-utilities) (73). The matching between the reported host strain in the PLSDB and the predicted host strain from the spacers database was tested at each taxonomic level, from the species level to the superkingdom level. For each plasmid with a match (a “matched plasmid” hereafter), the lowest taxonomic level at which the reported host and the predicted host match was recorded. The concordance for each taxonomic level is defined as the percentage of plasmids for which there was an agreement between the reported host and at least one of the predicted hosts at said taxonomic level.

### Host range prediction

For each matched plasmid, we identified the highest taxonomic level at which different hosts could be found. Using our predicted hosts, we assigned a host range grade based on the scale defined in (27), wherein plasmids for which all the hits differed only at the species level were assigned to Grade I; at the genus level to Grade II; at the family level to Grade III; at the order level to Grade IV; at the class level to Grade V; and at the phylum level to Grade VI. Plasmids were classified as mobile (MOB+) or non-mobile (MOB-) based on the presence or absence, respectively, of a MOB group, by using COPLA (74) with default parameters. Plasmids with only one hit or with several hits from the exact same host species were also assigned to Grade I.

### Network analysis of matched plasmids

A table containing a list of all the matched plasmids and the taxonomic lineage of their predicted hosts (18,475 records) was prepared. The table was imported to Cytoscape Ver. 3.9.1 (75), and a bipartite network was drawn using forced-directed layout with plasmids and bacterial families defined as nodes and the predicted presence of a plasmid within a host from a given family defined as an edge between the plasmid and the corresponding family. Families were colored according to the phylum to which they belong. Isolated families, namely, families that were not connected to other families via plasmids and that contain less than 10 plasmids, were removed from visualization for the sake of clarity.

### Network analysis of PTUs

The accession numbers from the list of the matched plasmids were matched to the accession numbers from the list of PTUs taken from supplementary table 2 in Redondo-Salvo et al. (27). All plasmids that mapped to the same PTU were collapsed into a single record, and the hosts of those plasmids were combined to produce a list of PTUs and their potential hosts. The table was imported to Cytoscape Ver. 3.9.1 (75) and a bipartite network was drawn as described above for the matched plasmids, using forced-directed layout with PTUs and bacterial families defined as nodes and the predicted presence of a PTU within a host from a given family defined as an edge between the PTU and the corresponding family. Families were colored according to the phylum to which they belong.

### Characterization of the CRISPR targets

The full features list for all the matched plasmids was retrieved using the Entrez Programming Utilities (E-utilities). The location of each hit on the plasmid was compared with the features list, and whenever a hit was found to be within an annotated feature, the feature type, feature ID, feature name and feature GO function were recorded. The relative frequency of each feature ID was recorded, and the 250 most frequent features were subsequently manually categorized into five groups: 1. Function Unknown: This group encompasses features categorized as ‘hypothetical protein’ and various DUF (domain of unknown function). 2. Plasmid Backbone Genes: Features in this category include genes associated with plasmid specific functions: conjugation, plasmid replication control, plasmid partitioning, and plasmid-borne toxin-antitoxin systems. 3. Potential Backbone Genes: This group includes genes associated with functions that may be related to plasmids but lack specificity, such as general DNA-modifying enzymes, DNA methylation, and restriction/anti-restriction functions. 4. General Functional Genes: Encompassing genes not necessarily associated with plasmids, this category includes metabolic genes, general ATPase, ribosomal RNA, and phage-associated genes. 5. Genes Associated with Transposons: This group comprises genes related to transposons, such as IS-associated integrase and transposase genes.

## ACKNOWLEDGMENTS

We thank Maria Pilar Garcillan Barcia and Tal Shay for fruitful discussion. This study was supported (in part) by grant no. 3-17700 from the Office of the Chief Scientist, Israel Ministry of Health, as a part of the MAPMAR project (a part of the ERA-NET Cofund AquaticPollutants project). LA is the recipient of a Hi-Tech, Bio-Tech, and Chemo-tech fellowship of Ben-Gurion University of the Negev.

